# Systematic Investigation of Machine Learning on Limited Data: A Study on Predicting Protein-Protein Binding Strength

**DOI:** 10.1101/2023.10.03.560786

**Authors:** Feifan Zheng, Xin Jiang, Yuhao Wen, Yan Yang, Minghui Li

**Affiliations:** Department of Bioinformatics, School of Biology and Basic Medical Sciences, Soochow University, Suzhou 215123, China

## Abstract

The application of machine learning techniques in biological research, especially when dealing with limited data availability, poses significant challenges. In this study, we leveraged advancements in method development for predicting protein-protein binding strength to conduct a systematic investigation into the application of machine learning on limited data. The binding strength, quantitatively measured as binding affinity, is vital for understanding the processes of recognition, association, and dysfunction that occur within protein complexes. By incorporating transfer learning, integrating domain knowledge, and employing both deep learning and traditional machine learning algorithms, we mitigate the impact of data limitations and make significant advancements in predicting protein-protein binding affinity. In particular, we developed over 20 models, ultimately selecting three representative best-performing ones that belong to distinct categories. The first model is structure-based, consisting of a random forest regression and thirteen handcrafted features. The second model is sequence-based, employing an architecture that combines transferred embedding features with a multilayer perceptron. Finally, we created an ensemble model by averaging the predictions of the two aforementioned models. The comparison with other predictors on three independent datasets confirmed the significant improvements achieved by our models in predicting protein-protein binding affinity. The source codes for these three models are available at https://github.com/minghuilab/BindPPI.

## Introduction

Currently, deep learning techniques have demonstrated impressive predictive capabilities when applied to large datasets^1,2^. However, many biomedical problems suffer from a scarcity of experimental data, making it imperative to explore how existing techniques can be utilized to achieve enhanced predictive accuracy^3^. In previous work, we have developed a set of data-driven machine learning methods to evaluate the impact of missense mutations on protein stability^4^ and their interactions with other molecules^5–10^. These methods employed traditional machine learning algorithms alongside handcrafted features, making them stand out as one of the widely recognized tools^11–13^. Additionally, traditional machine learning algorithms provide a direct measure of feature importance, enhancing the interpretability of the model outcomes. However, this approach has faced a bottleneck, leading to a lack of substantial improvement in predictive performance in recent years^14^. Despite the introduction of deep learning approaches in this field, their performance improvement has been severely hampered^15–17^, primarily due to the limited availability of training data. Continued efforts are required to overcome the challenges posed by limited data in machine learning. In this study, we leveraged the advancements in protein-protein binding affinity methods to conduct an in-depth investigation of the application of machine learning on datasets with limited data availability.

Protein-protein interactions (PPIs) play a fundamental role in various mechanisms of protein biological functions^18,19^ and become attractive targets for therapeutic intervention^20,21^. Numerous interactions between intracellular proteins are in principle possible but only a fraction of putative complexes is formed and functionally relevant Thus, the determination and characterization of structures and binding strength of protein-protein associations can gain significant insight into mechanisms in biological processes for disease research, such as prominent disorders of cancer and degenerative diseases associated with aberrant PPIs^22^. In therapy, one goal is to design new synthetic protein-protein complexes with the desired function, such as optimized antibody-antigen interactions with strong binding^23^. Thus, the characterization of PPIs in terms of their binding strength is highly relevant to the design of new and improved therapeutics^24,25^. Binding affinity is a quantitative measure of the strength of the interaction between two or more molecules that bind reversibly. Experimental techniques, such as surface plasmon resonance^26^, isothermal titration calorimetry^27^, and fluorescence resonance energy transfer^28^, for measuring binding affinity require expensive experimental setup and are time-consuming. For this reason, developing computational methods to predict binding affinity is increasingly important, which can help evaluate and understand the significance of putative protein-protein interactions^29^, the discovery of protein therapeutics^25^, *de novo* interface design^30^, etc.

The computational prediction of binding affinity has a long history and various methods have been proposed throughout the years, varying dramatically in terms of accuracy, computational cost, and physical plausibility^31^. Sophisticated approaches, such as free energy perturbation (FEP)^32^ and thermodynamic integration (TI)^33^, and end-point methods, like molecular mechanics Poisson–Boltzmann surface area (MMPBSA)^34,35^, possess a relatively high level of accuracy in principle. However, these methods that employ extensive molecular dynamics or Monte Carlo conformational searches are computationally intensive and have a limited scope of application. Alternative, simplified empirical energy functions have been proposed to significantly reduce computational costs. One such method is statistical potentials, which uses the observed relative positions of atoms or residues in experimental structures to infer a potential of mean force^36,37^. Another approach that has gained increasing popularity over the past decade is machine learning, where energy functions are determined through regression against experimentally measured binding affinities^16,38–46^. However, the prediction accuracy of currently available methods remains limited. Hence, it is essential to keep putting effort into developing accurate and reliable methods to tackle the challenge of predicting protein-protein binding affinity.

In this research, by exploring transfer learning, integrating domain knowledge, and utilizing both deep learning and traditional machine learning algorithms, we mitigate the impact of data limitations and make significant advancements in predicting protein-protein binding affinity. Specially, we compiled a dataset of 802 protein-protein complexes with reliable experimental measurements of binding affinities and complex structures. We developed more than 20 predictive models belonging to four categories: structure-based models with handcrafted features, sequence-based models with transferred embedding features, ensemble models composed of structure-based and sequence-based models, and structure-sequence models with a combination of handcrafted and embedding features. Among these models, three were selected with the best performance representing the three categories. The structure-based model is composed of a random forest regression and thirteen carefully selected handcrafted features derived from the complex structures. Our sequence-based model consists of a multilayer perceptron and average pooling of embedded features extracted from ESM-2^47^. By combining these two models, we obtained an ensemble model that outperforms each individual model. To validate the advancements achieved by our approaches, we compared the performance of our methods with other previously published predictors using three independent datasets. The results demonstrate the significant improvements achieved by our models.

## Methods

### Experimental datasets used for parameterizing our methods

The training dataset was compiled from four databases/datasets: Protein-Protein Binding Affinity Benchmark version 2 (PPBABv2)^48^, SKEMPI 2.0^49^, PROXiMATE^50^, and PDBbind version 2020^51^. These databases contain experimentally measured binding affinities and three-dimensional (3D) complex structures for protein-protein interactions (PPIs). The binding affinity was calculated using the equation Δ*G_exp_* = *RTln*(*K_D_*) = *RTln*(*K_i_*) = *RTln*(*IC*_’50_). Specifically, we collected 179, 348, 118, and 1306 protein-protein interactions from PPBABv2, SKEMPI 2.0, PROXiMATE, and PDBbind, respectively. To ensure data quality, we removed entries with ambiguous affinity values. Then, we merged all these four datasets into a combined dataset. In cases where the same PPI entry had multiple affinity values, we first filtered out entries with standard deviations of multiple affinity values larger than 1.0 kcal mol^-1^. Then, we selected only one affinity value based on the following criteria: (i) The priority was given to values that had been measured by more than one experimental technique or study, indicating a higher frequency of occurrence; (ii) We further prioritized values measured using surface plasmon resonance or isothermal titration calorimetry, as these methods are considered more reliable and accurate. If the above criteria still resulted in multiple values for a PPI entry, we calculated the average value. As a result, we obtained a total of 1562 unique protein-protein interactions with a single experimentally determined binding affinity value and their corresponding 3D structure.

One of the primary objectives of this study is to establish a structure-based model. To ensure the highest possible resemblance between the 3D complex structures employed in constructing the theoretical model and the proteins utilized for measuring binding affinity, we applied the following criteria to exclude certain complexes: (a) Protein-peptide and peptide-peptide complexes were removed if a chain had fewer than 50 amino acids, defining it as a peptide (427 complexes removed); (b) Complexes with metal coordination sites or containing modified/unknown/missing residues at the protein-protein binding interface were removed (225 complexes removed). The interface residues were defined as those with inter-atomic distances less than 6 Å between any heavy atoms of the interacting protein partners. The removed 652 complexes were used to compile the independent test sets (see details in the next section). In the final step, a total of 108 multimers were removed and set aside to serve as one of the test sets. Our training set, referred to as S802, comprised a meticulously selected 802 heterodimers. For more detailed information about the dataset, please refer to Fig. 1 and Supplementary Table 1.

**Figure 1.**
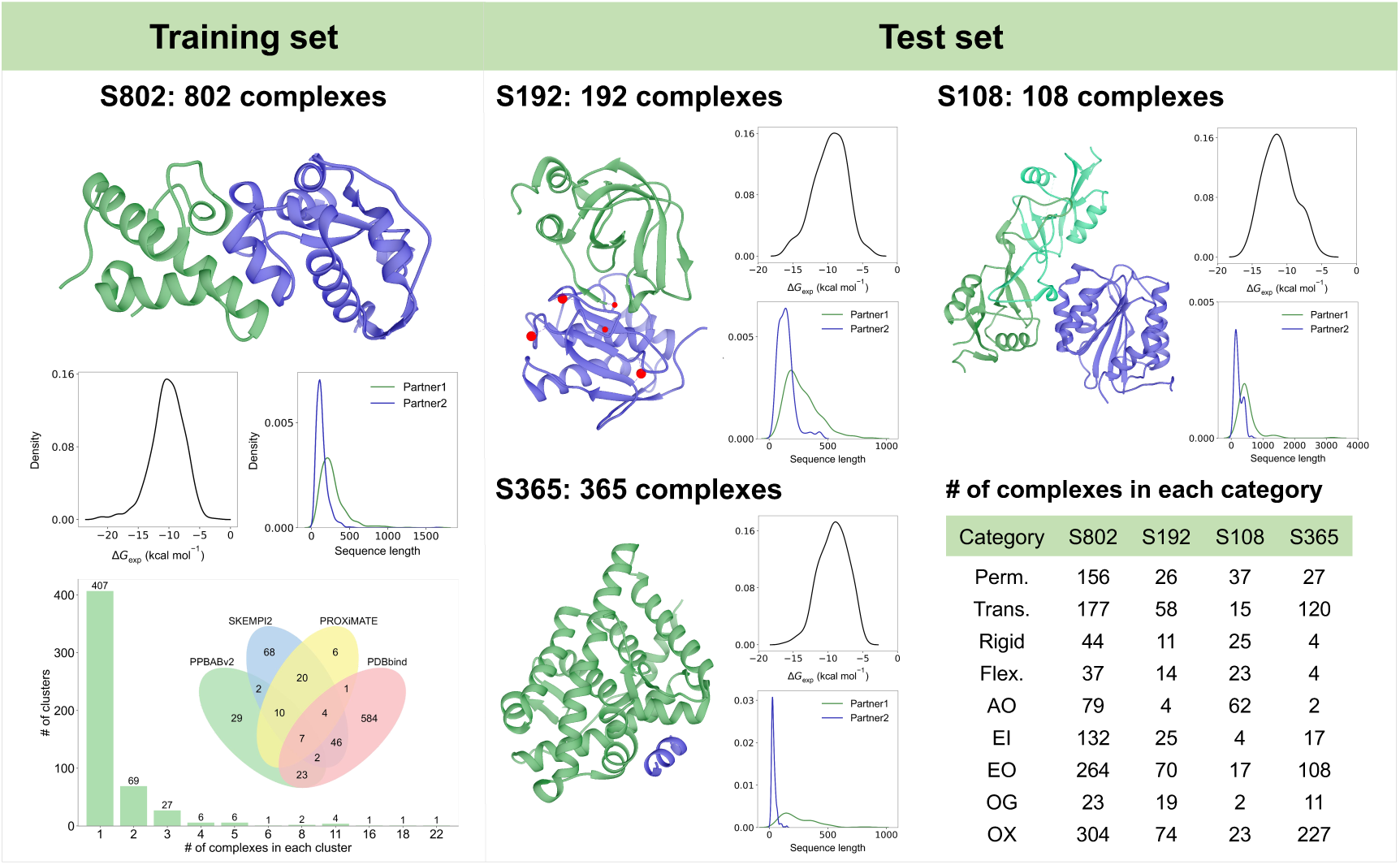
Overview of the data sets used. S802: visualization of protein-protein heterodimer complex structure, the distribution of experimental binding affinity and sequence length for each interaction partner, the four sources compiled from, and the number of clusters based on sequence similarity analysis; S192: visualization of heterodimer having metal coordination sites or modified/missing residues at the binding interface; S108: visualization of multimer. S365: visualization of protein-peptide complex structure, a chain with less than 50 amino acids were defined as peptide. Statistics of different classification of complexes. See Table S1 for more information.

### Experimental datasets used for testing

Initially, we compiled two test sets using the removed 652 complexes during the construction of the training set. We excluded specific types of complexes, such as peptide-peptide complexes, complexes with unknown residues at the binding interface, and protein-protein complexes with peptides at the binding interface (A peptide was defined at the binding interface if any of its residues belong to the interface residue set). Additionally, complexes with any individual missing interval at the binding interface greater than or equal to five amino acids were omitted. As a result, we retained 192 protein-protein heterodimer complexes (referred to as S192) and 365 protein-peptide complexes (referred to as S365) as our independent test sets. Secondly, we utilized the 108 multimers, which were previously removed during the construction of the training set (referred to as S108), to evaluate the performance of our method on multimers. Then, we took the following procedures to repair the complex structures. We converted modified residues to their corresponding standard residues. For missing segments at the binding interface, we employed the Modeller software^52^ to model them. Regarding complexes with metal coordination sites, we did not add the corresponding metals. For further information regarding the test sets, please refer to Fig. 1 and Table S1.

### Structure optimization

The complex 3D structures were obtained from the Protein Data Bank (PDB)^53^. Only assigned interaction partners were retained in the calculation, and missing heavy side-chain and hydrogen atoms were added by VMD program^54^ with the CHARMM36 force field parameters^55^. To optimize the structures, we tested several minimization procedures in the gas phase for all complexes to remove steric clashes or repair possible distorted geometries. These procedures included: (a) A 100-step energy minimization with restraints on the backbone atoms (the force constant is 5 kcal mol^-1^ Å^-2^); (b) A 2000-step energy minimization applying the same harmonic restraints as in (a); (c) A 2000-step minimization with restraints, followed by an unconstrained 5000-step minimization. The energy minimization was performed using the NAMD program (v 2.13)^56^ based on the topology file of CHARMM36 force field.

### Analysis of similarity among datasets

To assess the similarity between datasets, we employed three methods: sequence similarity analysis, structure similarity analysis, and their combined use. For sequence similarity analysis, we used the MMseqs2 software^57^ and set the sequence identity threshold to 50%, with the alignment covering at least 50% of both query and target sequences. To be considered similar, two complexes must have both protein chains with similar sequences. For structure similarity analysis, we used TM-align^58^ to perform structural alignments, focusing on the interface regions to compare the structure similarity of all protein-protein complexes. We generated a TM-score distance matrix encompassing all interface regions and conducted hierarchical clustering of the complexes using AgglomerativeClustering from the scikit-learn library^59^, employing a distance threshold of 0.3 (TM-score > 0.7) for clustering. The results of our analysis, as presented in Table S1b, revealed a substantial diversity among the complexes in the training set, highlighting the heterogeneous composition within the dataset. Furthermore, the similarity between the test sets and the training set was relatively low, indicating distinct characteristics between these sets.

### Construction of structure-based models with handcrafted features

Deep learning has received considerable attention in recent times due to its impressive performance when trained on large datasets. However, in scenarios with limited training data, traditional machine learning, combined with domain expert-identified features, continues to demonstrate comparable or even superior predictive capabilities to deep learning methods^11^. In our study, we initially identified and calculated numerous handcrafted features associated with protein-protein binding. An overview of all these features is provided in Table S2. It is worth noting that while certain features can be computed solely using the protein sequence, in our study, we classify them as structure-based features because we specifically utilized those features derived from interface or surface amino acids. Overall, we generated a list of more than a thousand descriptors.

In our previous studies^4–7^, the Random Forest (RF) algorithm has undergone rigorous validation and exhibited superior performance compared to other traditional machine learning algorithms. Moreover, RF offers a clear measure of feature importance and exhibits remarkable computational efficiency. Therefore, we first employed the RF algorithm to construct the structure-based models (Fig. 2a). Feature selection is an important step in traditional machine learning that enables us to identify and select the most relevant and informative features that can improve the model’s performance on unseen data while reducing the dimensionality of the input data and computational time. It can also help to increase the interpretability of the model’s results by focusing on the most important input features. In this study, we employed a modified scheme of the forward feature selection, which has been demonstrated to be effective in our previous studies^4–10^. This scheme is based on several principles, including the minimization of the number of features, maximization of their contribution to the model, ensuring low correlation among selected variables, and enhancing feature interpretability. When these principles are satisfied, the priority is determined by computational cost. As a result, a total of thirteen features from ten categories were selected. The selected features are described below, and their respective contributions to the model (referred to as RF_{13}) are presented in Table S3.

- *SA^p^*^1^ and *SA^p^*^2^ are the solvent accessible surface areas (SASA) of unbound partners (p1: partner 1 and p2: partner 2), which are calculated using the CORMAN module of CHARMM^60^.
- 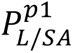 and 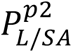 are the ratios of sequence lengths and solvent accessible surface areas for two unbound partners, which measure how tightly the protein structure is packed.
- 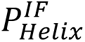 and 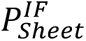 represent the percentages of helices and sheets at the binding interface, respectively, which are calculated by dividing the number of residues assigned in helix/sheet conformation by the total number of interface residues. The secondary structure elements are assigned using the DSSP program^61,62^.
- 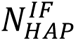 is the number of interactive heavy atom pairs between two partners. Two heavy atoms are considered interactive if their distance is within 6 Å.
- 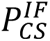 stands for the ratio of the number of conserved residues at the binding interface to the total number of interface residues. A residue is considered conserved if the score for a residue mutated to alanine, as calculated by PROVEAN^63^, is no more than −2.5.
- Δ*E_elec_* is the electrostatic interaction between two interacting partners, which is calculated as the difference in electrostatic energies between a bound complex and each interacting partner. The calculation is performed using the ENERGY module of CHARMM^60^.
- 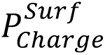 represents the percentage of charged amino acids on the surface of complex. It is calculated by dividing the number of surface charged residues by the total number of surface residues. A surface residue is defined by a SASA ratio greater than 0.2 between the residue in the complex and the extended tripeptide^64^. The SASA values for the residue in the extended tripeptide and complex are obtained from^65^ and calculated using DSSP program^61^, respectively.
- 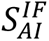 is sum of the amphiphilicity index of amino acids located at the binding interface. The amphiphilicity index of amino acids is obtained from Amino Acid Index Database^66^ with identifier MITS020101^67^.
- 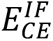 is the inter-protein contact energy calculated using atom-atom (AACE167) statistical contact potentials derived from the Potts model implemented in iPot program^68^. The contact distance cutoff is *d*_max_ = 10 Å, and the sequence separation is *k*_*min*_ = 5.
- *V_cavity_* represents the number of water molecules that can be accommodated in the cavities of the complexes, calculated using the McVol program^69^. Cavities are defined as empty spaces with sufficient volume to accommodate a water molecule.

**Figure 2.**
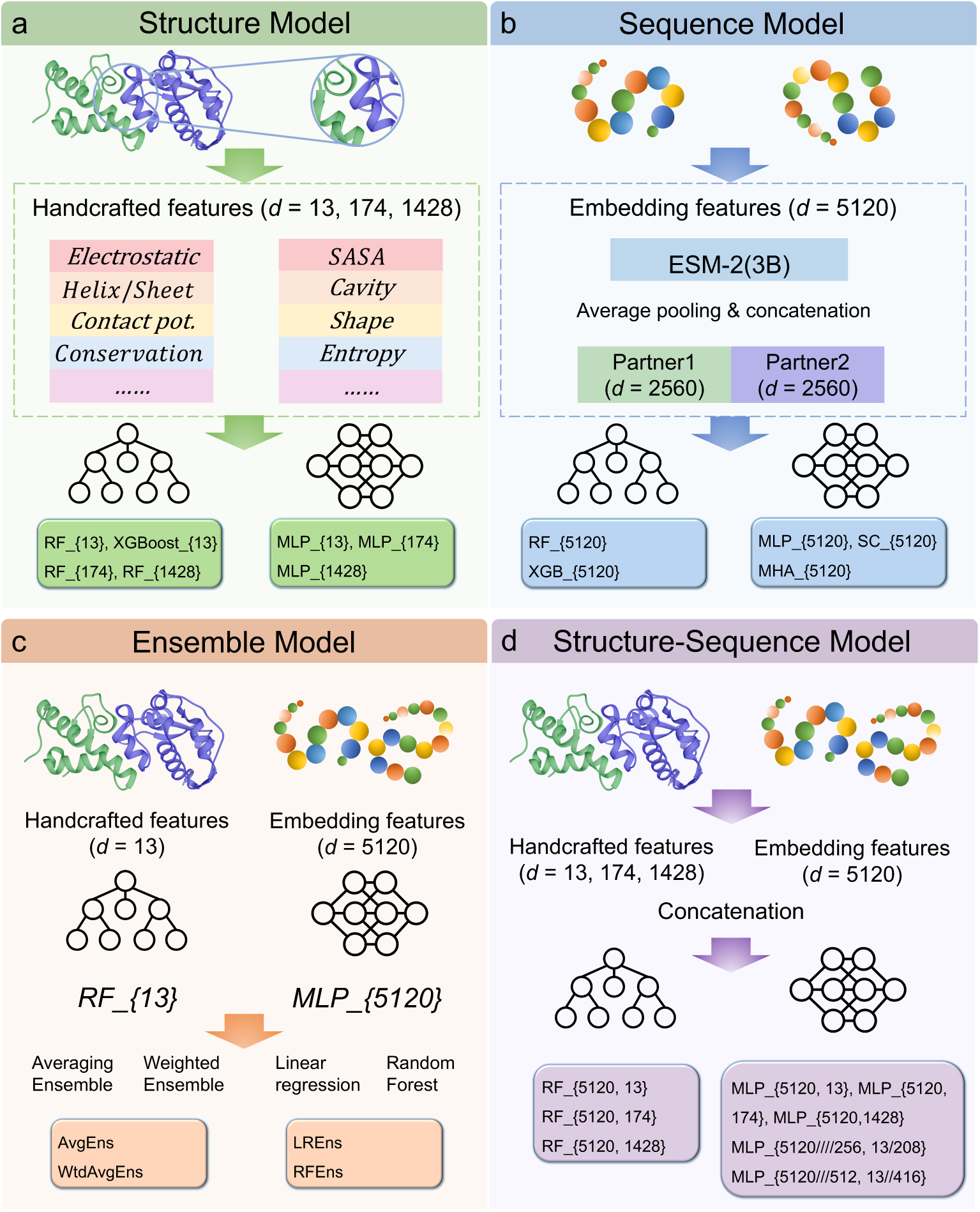
Overview of the framework, consisting of four types of models. (a) Structure-based models with handcrafted features, which include a comprehensive set of physicochemical, evolutionary, sequence, and structural features for in-depth feature representation. (b) Sequence-based models with embedding features, which use a fixed-length concatenated vector representation for each complex (*d* = 5120) obtained from a large-scale pretrained language model, ESM-2(3B). (c) Ensemble models, combining the best-performing structure and sequence models obtained from (a) and (b). (d) Structure-sequence models, integrating a combination of raw sequence embedding features and handcrafted structure features. See Table S4 for the definition of models.

In order to conduct a comparison with the RF_{13} model, we utilized a different traditional machine learning algorithm, eXtreme Gradient Boosting (XGBoost), and experimented with a deep learning multilayer perceptron (MLP) neural network that incorporated the handcrafted features. Additionally, we conducted a thorough evaluation to assess the importance of feature selection, analyzing not only the 13 selected features but also all available features (Table S4 provides a detailed description of all models).

### Construction of sequence-based models with transferred embedding features

For each protein sequence with a length of *L*, we extracted amino acid-level embeddings from the ESM-2(3B) model^47^, a large-scale pretrained language model based on the BERT transformer architecture. The ESM (Evolutionary Scale Modeling) models were trained to predict masked amino acids using the surrounding amino acids in the sequence. The ESM-2(3B) model consists of 36 transformer layers, containing 3 billion parameters, and was trained on over 60 million protein sequences. For each amino acid, the ESM-2(3B) model outputs a feature vector of dimension *d* = 2560. To represent a protein, we computed the average of all amino acid embeddings over the *L* amino acids using average pooling. This process resulted in a fixed-length vector representation of *d* = 2560 for each protein. For cases where one interaction partner contains multiple chains, we first average each individual chain and then calculate the average for all chains within the partner. Subsequently, we concatenated the two protein-level embeddings to obtain a 5120-dimensional feature vector for a protein-protein complex. This concatenated feature vector was used as the input for the sequence-based models in our study (Fig. 2b and Fig. S1a).

Initially, we constructed the sequence-based predictive models using the Random Forest and eXtreme Gradient Boosting algorithms (Table S4). Subsequently, we employed a multilayer perceptron (MLP) neural network to develop our predictive model (refer to Fig. S1a). The MLP architecture consists of multiple fully connected hidden layers followed by an output layer, with a rectified linear unit (ReLU) activation function applied after each hidden layer. The mean squared error (MSE) was used as the loss function to assess the model’s performance. To determine the optimal number of epochs, we incorporated an early stopping technique. We set a patience value of 20, considering an epoch improved only if its validation MSE loss surpassed the previous loss by a tolerance of 0. To ensure the robustness of our model, we conducted 10 iterations of 5-fold cross-validation. The selection of the optimal hyperparameter combination for each cross-validation was based on the Pearson correlation coefficient calculated on the validation set. Instead of refitting the model with the entire training set, we used the mean of 50 models for prediction. The results of our experiments confirmed that this architecture achieved the highest performance among the evaluated models and was consequently selected as our final sequence-based model (named as MLP_{5120}).

In addition to the aforementioned architectures, we enhanced our feature processing procedure by incorporating a multi-head attention network and skip connections. One architecture, shown in Fig. S1b, utilized two fully connected layers to capture patterns within individual proteins. Another architecture, illustrated in Fig. S1c, involved using a multi-head attention (MHA) layer to obtain fused embeddings for each protein, which could capture the interactions between proteins. The transformed embeddings from these layers were then concatenated. To further facilitate information flow, skip connections were introduced in these two architectures. Finally, both models incorporated an MLP architecture to further process and utilize the embeddings obtained from the skip connections. By incorporating these techniques, including multi-head attention, skip connections, and MLPs, we aimed to enhance the representation of individual proteins and protein complexes, capturing their complex relationships and improving the overall model performance. However, It is worth noting that these architectures did not outperform MLP_{5120}.

### Construction of ensemble models combining structure and sequence models

Based on the correlation analysis of the prediction results obtained from the sequence models and the structural models (Fig. S2), we propose that combining these two types of models can further enhance the predictive performance (refer to Fig. 2c and Table S4). To implement the ensemble, we explored three distinct approaches: averaging, weighted averaging, and stack-based methods. In the weighted average approach, the weights were determined based on the Pearson correlation coefficient values of each model obtained from 5-fold cross-validation. To ensure that the scaling of predicted values was unaffected, we normalized the weights of each model in each weighted ensemble combination. For the stacking method, the predicted results from the individual models were utilized as input features to train a meta-model. The meta-regressor was constructed using two algorithms: linear regression and random forest. The results confirmed that the average and weighted ensemble approaches achieved the highest performance, and consequently, the simplest average ensemble was selected as our final ensemble model.

### Construction of structure-sequence models combining handcrafted and embedding features

Finally, we integrated a combination of raw sequence-based embedding features and structure-based handcrafted features to build the predictive models using the MLP and RF algorithms, respectively (Fig. 2d and Table S4). First, we directly concatenated the 5120-dimensional sequence features with the handcrafted features, which have dimensions of 13,174, and 1428, as inputs to construct MLP and RF models. However, due to the substantial dimensionality gap between the sequence features (*d* = 5120) and the 13 selected one-dimensional structure features, we adopted a two-step method to address this issue, as depicted in Fig. S1d. In the first step, we independently adjusted the dimensionality of the structural and sequence features. For the structural features, we increased their dimensionality by employing one or two layers, resulting in either 16 or 32 dimensions for each feature. Consequently, the dimensionality of the 13 structural features increased to either 208 or 416 dimensions. Simultaneously, we reduced the dimensionality of the 5120-dimensional embedding to 512 or 256 dimensions using three or four layers, respectively, to match the dimensionality level of the structural features. In the second step, we concatenated the up-sampled and down-sampled features, which were now at compatible dimensional levels, and utilized them as inputs to perform the MLP architecture.

### Hyperparameter turning

The hyperparameters of the RF and XGBoost models were optimized through a grid search approach within a predefined hyperparameter search space. The complete list of hyperparameters can be found in Supplementary Table 5. The optimal combination of hyperparameters was chosen based on the average Pearson correlation coefficient obtained from 5-fold cross-validation. The final RF and XGBoost models were trained on the entire training set using these selected hyperparameters.

To improve computational efficiency, a sequential search strategy was employed for hyperparameter selection in MLP models. The hyperparameters were determined sequentially, beginning with the number of hidden layers, followed by the learning rate, batch size, hidden layer dimension, and weight decay, as indicated in Table S5. The order of selection was based on the relative importance of each hyperparameter’s impact on the model’s performance. The Pearson correlation coefficient calculated on the validation set was used to choose the optimal combination of hyperparameters for each cross-validation. Instead of retraining the model with the entire training set, the mean of 50 models (10 repetitions × 5 folds) was used for prediction, enabling a more stable and reliable estimation of model performance.

### Classification of protein-protein interactions

In this research, protein-protein interactions were categorized into three distinct classes. The first classification was based on the strength of the interaction: Permanent interactions were identified by an interaction strength with a Δ*G_exp_* value of ≤ −12.27 kcal mol^-1^, while transient interactions were defined by an interaction strength with a Δ*G_ext_* value of ≥ −8.18 kcal mol^-1^.^70,71^ The second classification was based on the flexibility of the complexes, dividing them into rigid-body and flexible complexes.

Rigid-body complexes were characterized by an interface C-alpha root-mean-square deviation (I-RMSD) value of ≤ 1.0 Å, whereas flexible complexes exhibited an I-RMSD value of > 1.0 Å.^44^ To calculate the I-RMSD, the unbound components were superimposed onto their bound complexes, considering the C-alpha atoms of the interface residues. Only a subset of entries possesses 3D structures of unbound components. Fig. 1 and Table S1c present the number of complexes falling into each category.

The third classification was performed based on the functional categorization of protein-protein complexes. Among the four data resources, only the Protein-Protein Binding Affinity Benchmark provided functional classification for the complexes, with categories such as Enzyme-Inhibitor and Antibody-Antigen. To extend the functional classification to our entire dataset, we performed the following steps to categorize complexes into six functional classes: AN (Antigen-Nanobody): complexes where at least one chain can be found in the SAbDab-nano database^72^; AA (Antibody-Antigen): complexes where at least one chain can be found in the SAbDab database^73^; As a result of the limited number of instances in the AA and AN categories, we combined them into a single category named AO. EI (Enzyme-Inhibitor): one of the proteins in the complex has an EC number obtained from PDBe (Protein Data Bank in Europe) or PDB, and the other protein is annotated with the term “inhibit” in Pfam, InterPro, SCOP, or CATH databases integrated within PDBe^74^; EO (Enzyme-Others): complexes where any chain is annotated with an EC number, except for those falling under the EI classification; OG (G-protein-Others): complexes where any chain has annotations containing the term “G protein” in the molecule function of GO^74^, as well as annotations containing the terms “G protein” or “GTPase activity” in Pfam, InterPro, SCOP or CATH databases; OX (Others-miscellaneous): all remaining complexes that did not fall into any of the aforementioned functional categories.

### Comparison with other methods

We performed a comprehensive comparison between our models and eight other state-of-the-art methods used to calculate the binding energy, including PRODIGY^44,45^, PPI-Affinity^38^, PPA_Pred2^16^, Minpredictor^42^, ISLAND^43^, FoldX^75^, Rosetta^76^, and MMPBSA^8,77^. PPA_Pred2 and ISLAND are sequence-based approaches, while the rest are structure-based methods. Among these methods, PRODIGY, PPI-Affinity, PPA_Pred2, Minpredictor, and ISLAND have been trained on protein-protein binding affinity data. The number of overlapping complexes between their training sets and our datasets is provided in Table S6. It is worth mentioning that except for PRODIGY, all the other four machine learning methods are limited to calculating the binding energy of dimeric complexes.

The three methods, FoldX, Rosetta, and MMPBSA, are commonly used for calculating absolute energy in protein systems. The AnalyseComplex module of FoldX was used to calculate the binding energy. For Rosetta, the binding energy is calculated using the ddG Mover in RosettaScripts with the beta_nov16 score function. The resulting energy values are reported in Rosetta Energy Unit (REU), which are correlated with kcal mol^-1^.^78^ MMPBSA combines molecular mechanical energies with the Poisson−Boltzmann continuum representation of the solvent, which was expressed as the sum of van der Waals interaction energy, polar solvation energy, and nonpolar solvation energy. For further information regarding the specific details of our MMPBSA implementation, please refer to our previous study^8^.

Additionally, we assessed the predictive performance of ten docking scores obtained from the CCharPPI webserver^79^ in estimating binding affinity. Ten docking scores include: ZRANK^80^, ZRANK2^81^, RosettaDock^82^, pyDock^83^, FireDock^84^, FireDock (antibody-antigen energy function)^84^, FireDock (enzyme-inhibitor energy function)^84^, PISA^85^, PIE^86^, and SIPPER^87^.

### Performance evaluation

Pearson correlation coefficient (PCC) and root-mean-square error (RMSE) were used to quantify the agreement between experimentally-determined and predicted values of binding affinities. A two-tailed *t*-test was used to assess whether the correlation coefficient is statistically significant from zero. RMSE (kcal mol^-1^) is the standard deviation of the prediction errors, calculated by taking the square root of the average squared difference between predicted and experimental estimates. To evaluate the statistical significance in the difference of PCC between our models and other methods, we employed the Hittner2003 test^88^, which is used for comparing two correlation coefficients based on dependent groups. Furthermore, we compared the receiver operating characteristics (ROC) curves using the DeLong test^89^.

To assess the performance of the proposed approaches in distinguishing permanent or transient interactions from others, ROC analysis was conducted. The true positive rate (TPR) and false positive rate (FPR) were calculated as follows: TPR = TP/(TP+FN) and FPR = FP/(FP+TN), where TP represents true positives, TN denotes true negatives, FP signifies false positives, and FN stands for false negatives. To account for imbalances in the labeled dataset, the Matthews correlation coefficient (MCC) was also computed. The Hittner2003 test was performed using the *cocor* package^90^ in R, while the remaining evaluation metrics were implemented using the Python SciPy^91^ and Scikit-learn packages^59^.

## Results and Discussion

### Structure-based models with handcrafted features

In our study, a total of 1,428 handcrafted features related to protein-protein binding were identified and computed (an overview of all these features is provided in Table S2). Subsequently, we constructed seven structure-based models using those features using the Random Forest, eXtreme Gradient Boosting, and multilayer perceptron neural network algorithms, respectively (Fig. 2a and Table S4). The Pearson correlation coefficient (PCC) and root mean square error (RMSE) values between the predicted and experimentally-determined binding affinities across all datasets are shown in Table S7. Overall, the models built using the RF algorithm exhibited better performance compared to those built using MLP and XGBoost algorithms. Furthermore, for the three RF models, an increase in the number of features resulted in a decrease in performance on the training set, while there was no significant change in performance on the test sets. In contrast, the MLP model performed poorly when built using only 13 features; however, as the number of features increased, the model’s performance improved significantly. This implies that the MLP model has better adaptability to more comprehensive feature representations.

Hence, based on the model performance, interpretability, and computational efficience, RF_{13}, composed of a random forest regression and thirteen carefully selected handcrafted features derived from the complex structures, was selected as our final structure-based model. This model achieved superior performance across the majority of datasets in comparison to other models, as evidenced by higher PCC and lower RMSE values (Fig. 3 and Table S7). To optimize the hyperparameters of RF_{13}, we conducted 5-fold cross-validation to evaluate different combinations of decision trees and features considered when splitting a node. Through this process, the optimal settings for RF_{13} are 230 decision trees and 2 features for splitting a node. Additionally, we utilized 100-step energy minimized complex structures for feature calculation, as further increasing the number of minimization steps did not yield improvements in prediction performance (Fig. S3).

**Figure 3.**
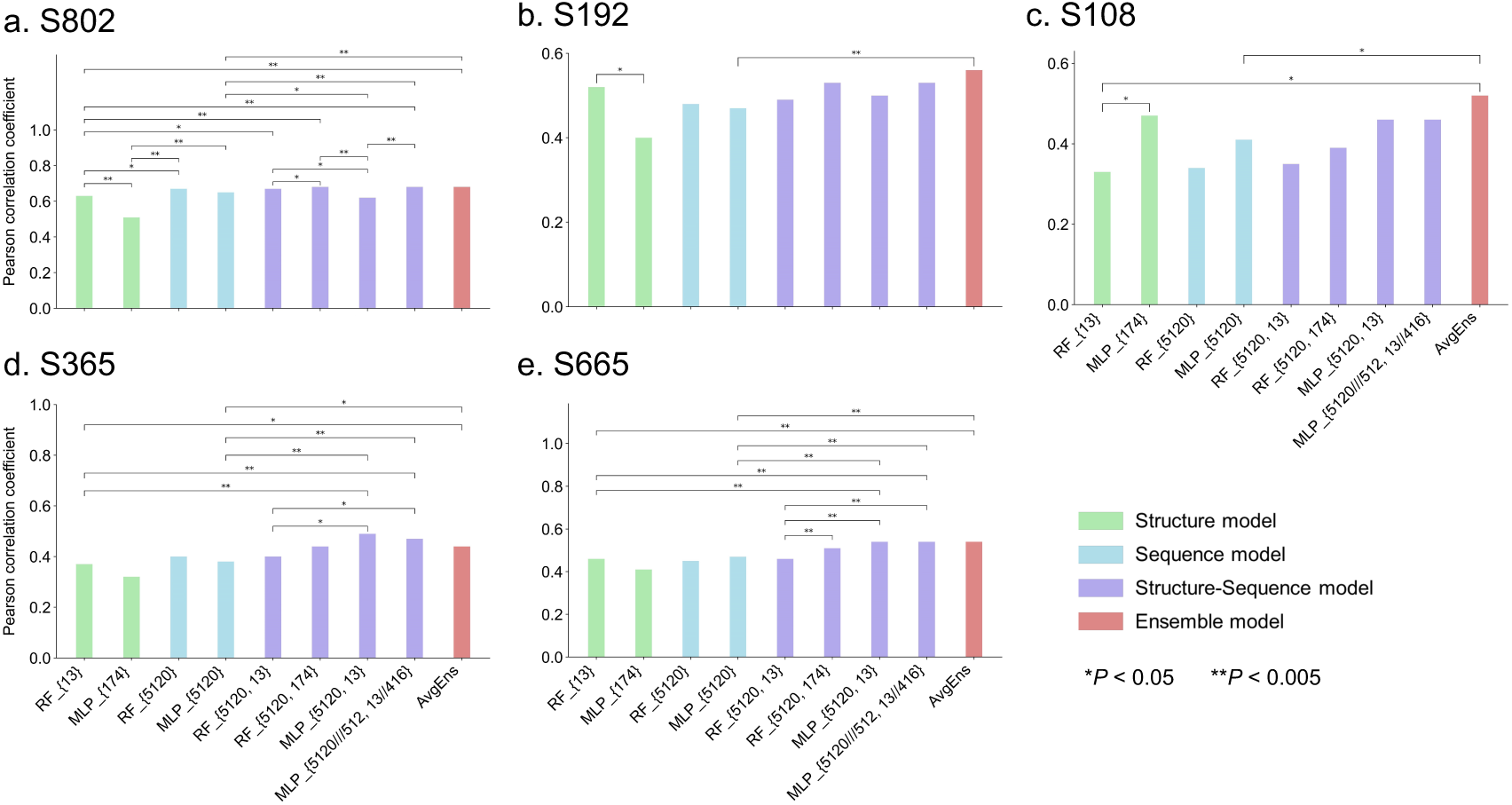
Pearson correlation coefficient between experimentally-determined and predicted values of binding affinities for nine models tested on five datasets. (a) S802, 5-fold cross-validation results are shown for S802. (b-d) three independent test sets of S192, S108, and S365. (e) S665, a combination of the three test sets. All correlation coefficients presented are significantly different from zero (*P* < 0.005, *t*-test). Significant comparisons were performed within four individual models and four structure-sequence models, as well as between the four structure-sequence models and RF_{13} and MLP_{5120}, respectively. The averaging ensemble model was also compared to RF_{13}, MLP_{5120}, and MLP_{5120///512, 13//416}, respectively. P values were calculated using the Hittner2003 test for comparing two correlation coefficients.

### Sequence-based models with transferred embedding features

Although AlphaFold and similar methods have greatly advanced our ability to predict the structures of individual protein monomers, predicting the structures of protein complexes remains a complex and challenging task^92^. Due to the limited availability of experimentally-determined protein complex structures, it is necessary to develop sequence-based models for our purpose of predicting protein-protein binding affinity. Here, we developed five models, including two traditional machine learning models and three deep learning models (Fig. 2b, Fig. S1a-c and Table S4). These models utilize the embeddings extracted from ESM-2(3B) as their input. Pre-trained models, trained on large-scale data, have learned complex feature representations of protein sequences and structures, and are widely used as features for numerous downstream tasks.

The performance evaluation based on PCC and RMSE values indicates that the RF model slightly outperforms the XGBoost model on the test sets (Table S7). During the exploration of deep learning architectures, it was observed that integrating multi-head attention and skip connections did not lead to a performance improvement beyond that of the simple MLP model. This suggests that the MLP architecture is already proficient in extracting pertinent information from the input features, making the added complexity of multi-head attention and skip connections unnecessary for our particular prediction task.

When comparing the best traditional machine learning model, RF_{5120}, with a deep learning model, MLP_{5120}, we observed that the PCC values were unable to differentiate between them (Fig. 3). Therefore, we proceeded to examine the distribution of predicted values and their performance on three different classification tasks. The results reveal that the RF model tends to produce more concentrated predictions within a limited range, while the MLP exhibits a broader distribution of predictions (Fig. S4). Upon assessing the performance across the three classification tasks, as depicted in Figure S5, it became evident that the RF model exhibits comparatively lower predictive capability than the MLP model in predicting flexibility complexes and functional classifications. This observation suggests that deep learning architectures, exemplified by the MLP model, possess a heightened ability to comprehend intricate patterns and representations from high-dimensional feature vectors compared to traditional machine learning techniques. In light of these findings, MLP_{5120} was designated as our final sequence model.

### Ensemble models combining structure and sequence models

As shown in Figure S2, the correlation between the prediction results of sequence and structural models is relatively low, as demonstrated by a PCC of 0.47 between RF_{13} and MLP_{5120} when tested on dataset of S665. We explored various combinations of the sequence and structural models and discovered that the averaging ensemble of the two top-performing models, RF_{13} and MLP_{5120}, yielded the top-level overall performance. This is understandable because the sequence models and the structural models themselves are highly correlated. The combined performance of RF_{13} and MLP_{5120} outperforms each individual model (Fig. 3). Furthermore, we explored weighted averaging and stack-based ensemble methods, but they did not exhibit higher performance compared to the simplest average ensemble (Table S7). Consequently, we selected the simplest average ensemble, referred to as AvgEns, as our final ensemble model. By doing so, we effectively exploit the individual strengths of each model while mitigating the limitations of their standalone predictions, resulting in a more robust and accurate overall predictive framework.

### Structure-sequence models combining handcrafted and embedding features

Finally, we employed a combination of raw sequence-based embedding features and structure-based handcrafted features to construct a total of eight predictive models, utilizing both MLP and RF algorithms, as depicted in Fig. 2d and Table S4. The evaluation results presented in Fig. 3 and Table S7 indicate that the incorporation of handcrafted features into MLP_{5120} does not result in a significant improvement in predictive performance on the test sets of S192 and S108. This observation holds true even when substantial efforts were made to narrow the dimensionality gap between the structural and sequence features. Similarly, integrating embedding features into the traditional RF models did not yield a notable enhancement in predictive performance across all test sets. Based on the results from S802 and S192, MLP_{5120///512, 13//416} was selected as the representative model within this category of models.

In summary, we have selected four models: RF_{13}, MLP_{5120}, AvgEns, and MLP_{5120///512, 13//416}, each representing distinct approaches in predicting protein-protein binding affinity. RF_{13} is a traditional machine learning model that employs 13 carefully selected structure-based handcrafted features, providing interpretability to the predictions. On the other hand, MLP_{5120} adopts a deep learning approach utilizing sequence-based transferred embedding features, offering advantages in terms of computational speed and independence from the 3D structure of the complex. Both models exhibit comparable performance, as demonstrated in Figure 3. The AvgEns model combines the predictions of RF_{13} and MLP_{5120}, leveraging the complementary strengths of both models and yielding superior predictive performance compared to individual models (Fig. 3 and Fig. 4). Lastly, MLP_{5120///512, 13//416} represents an attempt to integrate the raw sequence and structure-based features. However, the performance of this integrated model is lower than that of AvgEns. As a result, AvgEns was chosen as our final combination model for predicting protein-protein binding affinity.

**Figure 4.**
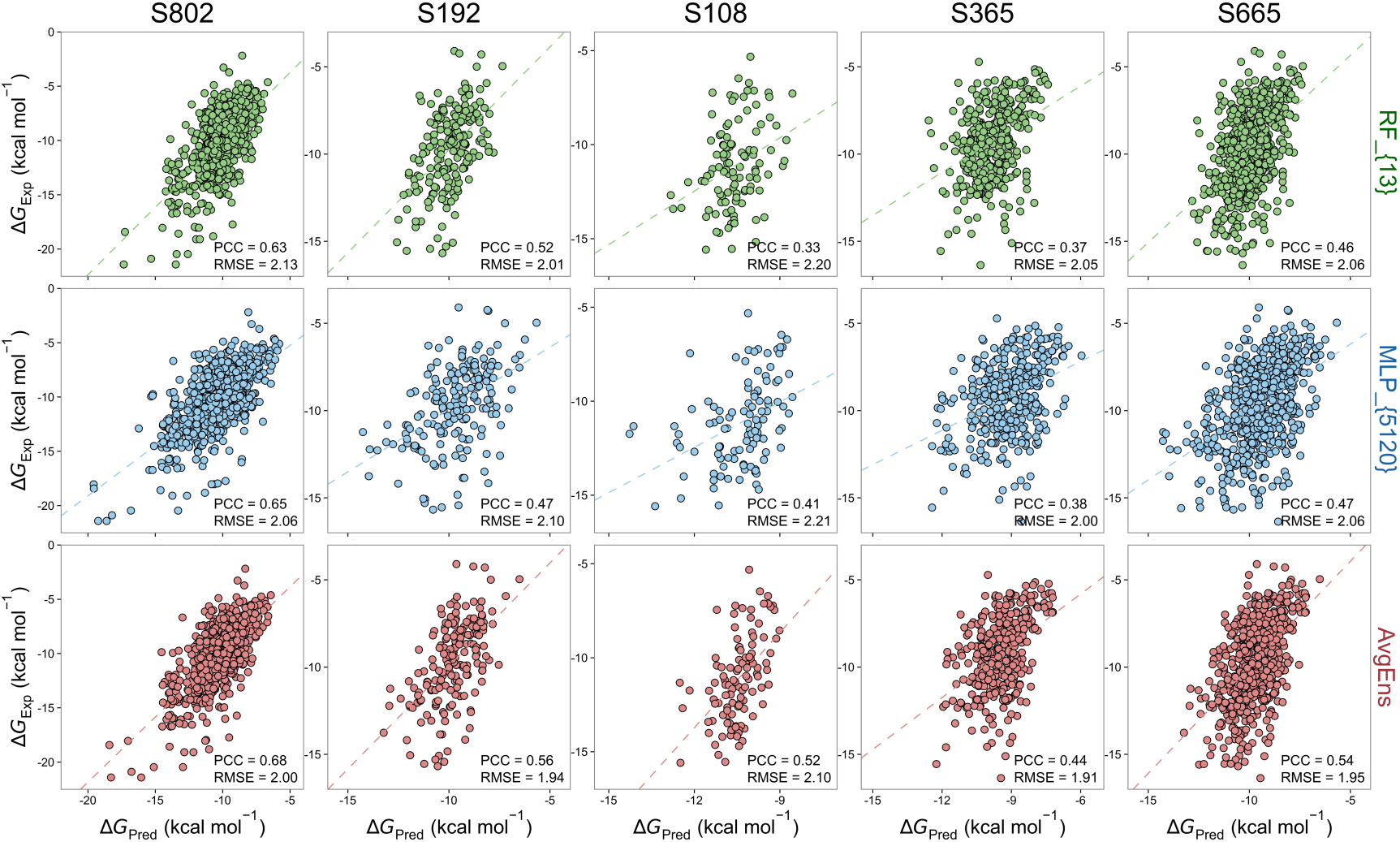
Performance of three selected representative methods, RF_{13}, MLP_{5120}, and AvgEns, on five datasets. 5-fold cross-validation results are shown for S802. All correlation coefficients are statistically significantly different from zero (*P* < 0.005, *t*-test). PCC: Pearson correlation coefficient, RMSE (kcal mol^-1^): root-mean-square error.

### Performance on protein-protein interaction classification

In this study, we conducted a classification analysis of protein-protein interactions based on three distinct characteristics: interaction strength, flexibility, and function of the complexes. We used S802 and S665 datasets to evaluate the performance, and the results are presented in Fig. 5 and Fig. S5. In the prediction of interaction strength, the ensemble model AvgEns outperforms each individual model significantly (*P* < 0.05, DeLong test), while RF_{13} and MLP_{5120} show comparable performance. Regarding the prediction of flexibility, our approaches demonstrate equal proficiency in predicting both rigid-body and flexible complexes. Specifically, RF_{13} exhibites higher PCC values for rigid-body complexes compared to MLP_{5120}, although the difference is not significant (*P* > 0.05, Hittner2003 test). Concerning functional classes, previous studies have shown that different types of functional complexes cannot be equally well predicted ^41,46^. Our three models demonstrate good performance on three types of complexes (EI, EO, and OX), with statistically significant PCC values. However, no statistically significant correlation was observed for OG complexes from S665. Predicting the binding affinity of complexes involving antibodies has always been challenging. Nevertheless, our sequence-based model has achieved significant PCC values in this context, indicating promising performance in predicting the binding strength of antibody complexes.

**Figure 5.**
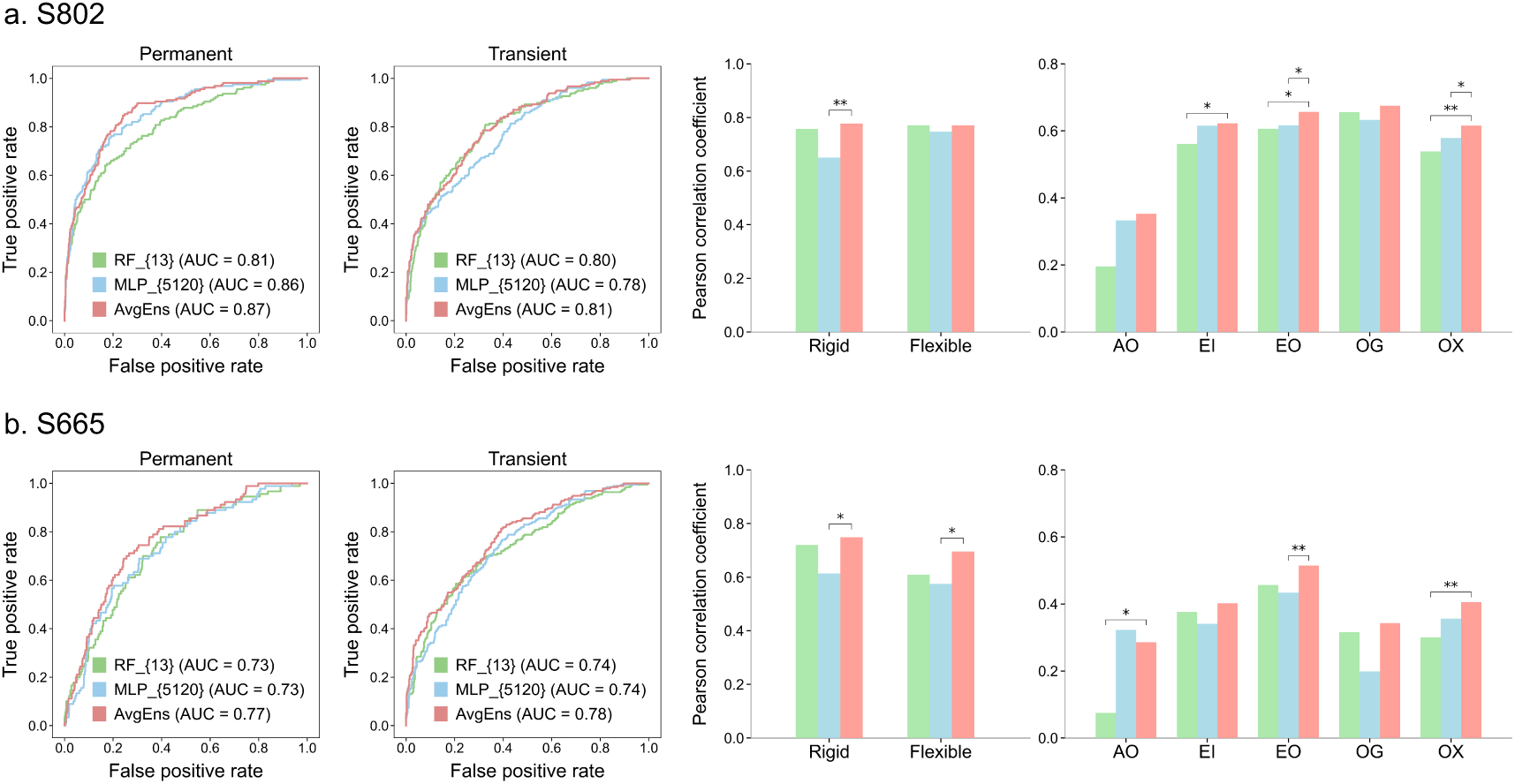
Performance of three methods, RF_{13}, MLP_{5120}, and AvgEns, was evaluated on datasets S802 (a) and S665 (b) for three distinct interaction classifications. Receiver operating characteristics curves for three approaches to distinguish permanent and transient protein-protein interactions from others. Pearson correlation coefficients for rigid-body and flexible complexes, and for five functional categorizations of complexes. The number of complexes in each category is provided in Fig. 1. P values were calculated using the Hittner2003 test for comparing two correlation coefficients (\**P* < 0.05 and \*\**P* < 0.005). The PCC values for RF_{13} applied on AO from both datasets do not have statistically significant difference from zero (*P* > 0.05, *t*-test). Additionally, the PCC values for all three methods applied on OG from S665 are not statistically significant either. The rest of PCC values are significantly different from zero (*P* < 0.05, *t*-test). See Supplementary Figure 5 for significant analysis of all PCC values.

### Comparative analysis with other approaches

Figure 6 and Table S8 present the performance comparison results of our three representative methods (RF_{13}, MLP_{5120}, and AvgEns) with five machine learning approaches trained on affinity data, three absolute energy calculation methods, and ten docking scores. Our methods consistently demonstrate superior performance compared to the other approaches across all datasets. Among the five machine learning approaches, PPI-Affinity shows relatively good performance with PCC values of 0.42 and 0.34 for S802 and S192, respectively. This performance may be attributed to the overlap between its training set with the S802 and S192 datasets (Table S6). The three absolute energy calculation methods (FoldX, MMPBSA, and Rosetta) are observed to be sensitive to different structure optimization approaches (Fig. S6). Hence, the highest PCC values for these methods on each dataset are reported. However, these methods exhibit limited predictive power, with non-significant or very low PCC values for S802 and S192 datasets. Additionally, the large RMSE values suggest that the predicted values from these methods cannot be directly interpreted as binding energy. As for the ten docking scores, they also exhibit very limited ability to predict affinity. This observation aligns with the understanding that the native conformation of the complex may not necessarily correspond to the one with the lowest binding energy, as the interaction is intended to generate a specific biological function rather than solely achieving high affinity.

**Figure 6.**
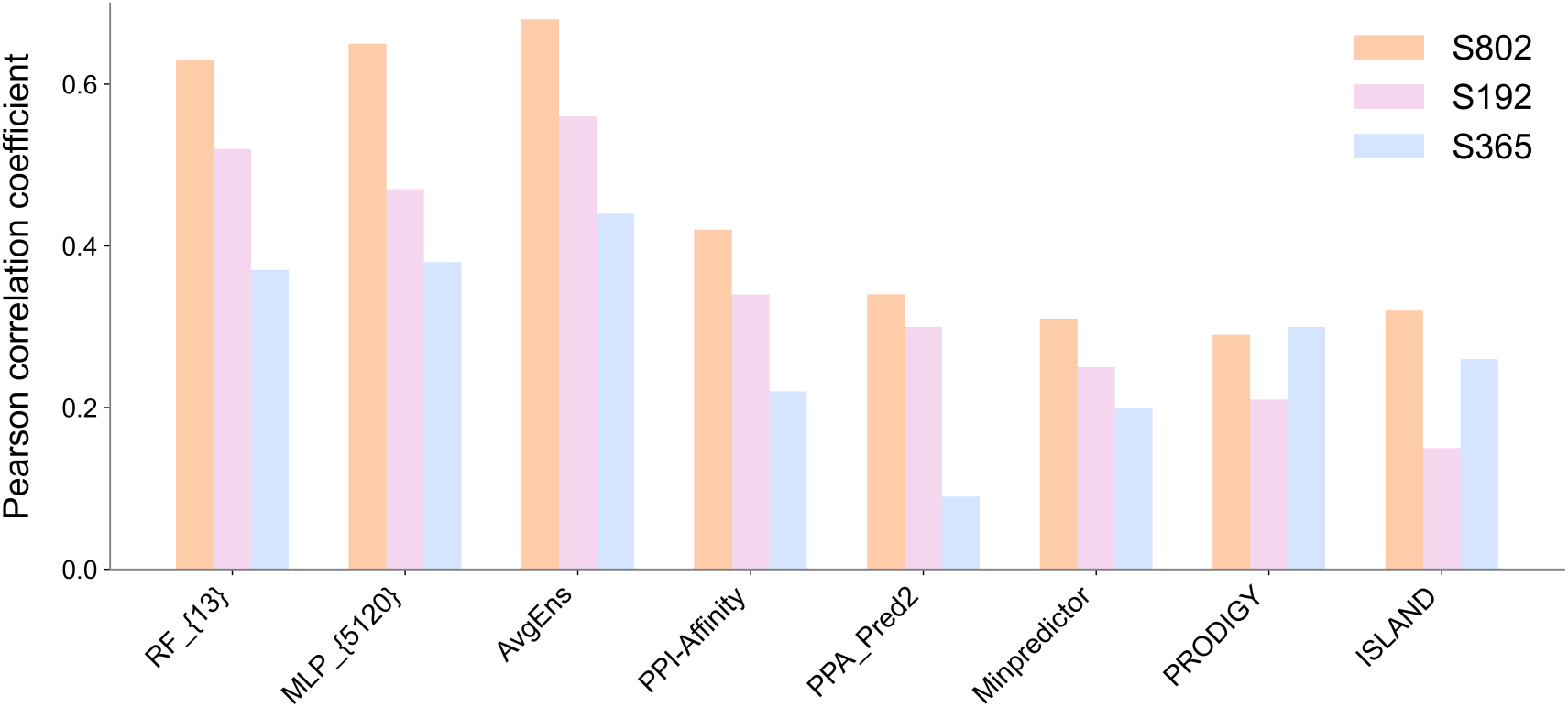
Comparison of methods’ performances. Pearson correlation coefficients are shown for our three representative methods and five machine learning approaches trained on binding affinity data. Among these five other approaches, only PRODIGY has the capability to predict multimers. Across all datasets, our method of AvgEns significantly outperform all the other approaches (*P* < 0.05, Hittner2003 test). The PCC value for PPA_Pred2 tested on S365 does not exhibit a statistically significant difference from zero (*P* > 0.05, *t*-test). For more detailed results, refer to Table S8.

Overall, in this work, we explored a range of approaches that incorporate transfer learning, integrate domain knowledge, and employ a combination of deep learning and traditional machine learning algorithms. These strategies collectively mitigate the impact of data limitations and lead to significant advancements in predicting protein-protein binding affinity. Our study yields the following insights:

- The integration of features extracted from extensive pre-trained models, when combined with deep learning techniques, yields promising predictive performance.
- Traditional machine learning methods, when coupled with carefully curated structural features grounded in prior knowledge, remain a valuable choice.
- The synergy between models leveraging transferred embedding features and those incorporating handcrafted features results in improved performance.

In light of these discoveries, we have devised sequence-based and structure-based methodologies that effectively estimate protein-protein binding affinity. These approaches hold great potential for aiding protein engineering endeavors by offering valuable starting points, minimizing the risk of unsuccessful laboratory experiments, and facilitating the design and development of therapeutic proteins.

## DATA AND CODE AVAILABILITY

The compiled experimental datasets and computational results that support our findings are publicly available on GitHub at https://github.com/minghuilab/BindPPI. The source codes of our three representative models are also available at https://github.com/minghuilab/BindPPI. Our methods are named as “BindPPI”. Additional data files and codes that support the findings of this study is available from the corresponding authors upon request.

## ACKNOWLEDGMENTS

This work was supported by the National Natural Science Foundation of China [32070665] and the Priority Academic Program Development of Jiangsu Higher Education Institutions. The funders had no role in study design, data collection and analysis, decision to publish, or preparation of the manuscript.

## AUTHOR CONTRIBUTIONS

Conceptualization, M.L.; Methodology, F.Z., X.J., Y.W., and M.L.; Software, F.Z. and X.J.; Validation, F.Z., X.J., and M.L.; Formal Analysis, F.Z.; Investigation, F.Z., X.J., and M.L.; Data Curation, F.Z. and X.J.; Writing – Original Draft, M.L.; Writing – Review & Editing, M.L.; Visualization, F.Z., Y.Y., and M.L.; Supervision, M.L.; Project Administration, M.L.; Funding Acquisition, M.L.

