## Supplementary Materials for "Systematic Investigation of Machine Learning on Limited Data: A Study on Predicting Protein-Protein Binding Strength"

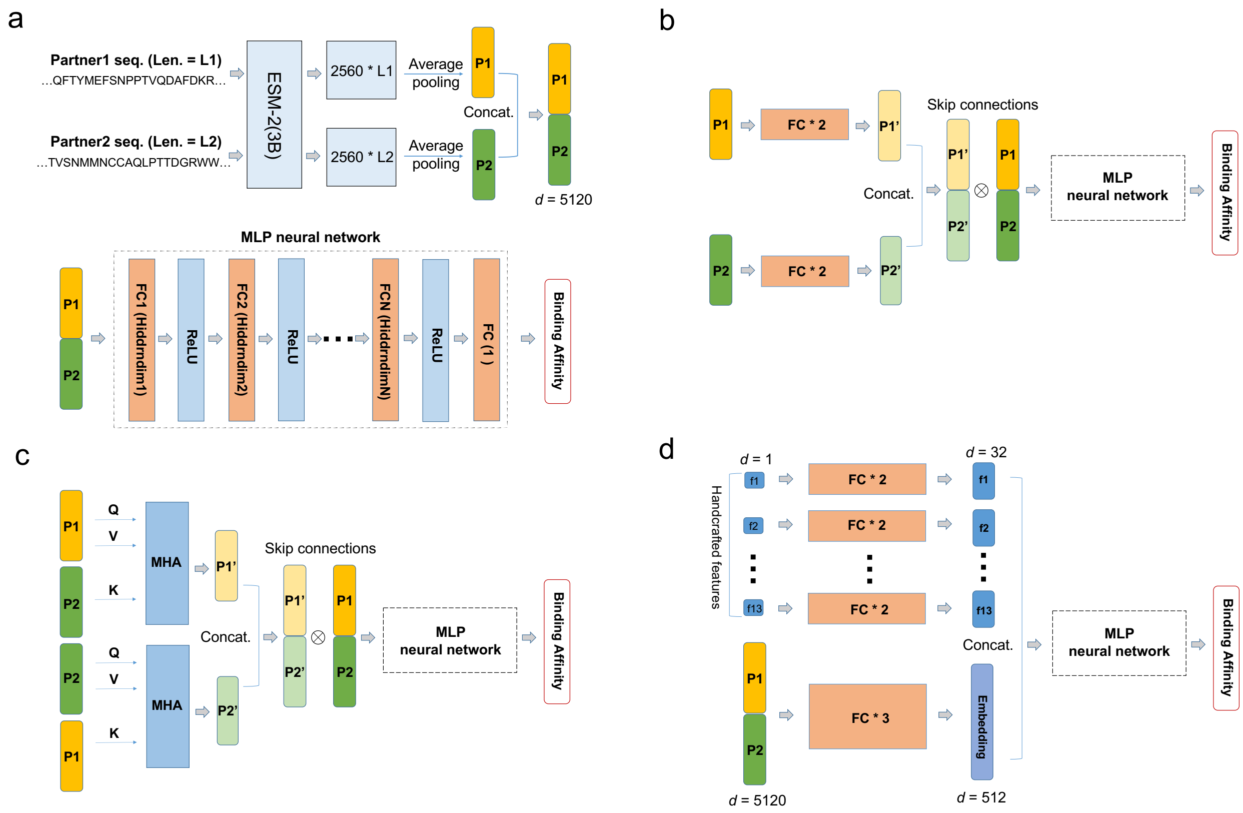


**Figure S1. Architectures of deep learning models.** (a) The multilayer perceptron (MLP) neural network. The architecture comprises multiple fully connected hidden layers followed by an output layer, with a ReLU activation function applied after each hidden layer. Inputs consist of a fixed-length concatenated vector representation ($d$ = 5120) obtained from ESM-2(3B). (b) The architecture employs two fully connected layers to capture patterns within individual proteins. (c) The architecture utilizes a multi-head attention (MHA) layer to obtain fused embeddings for each protein. The transformed embeddings from these layers are then concatenated to form complex-level embeddings. Skip connections were introduced in these two architectures (b and c). (d) The architecture for the MLP_{5120///512, 13//416} model involves increasing the dimensionality of each structural feature from 1 to 32 using two fully connected layers, and reducing the dimensionality of the 5120-dimensional complex embedding to 512 using three fully connected layers. Subsequently, the up-sampled and down-sampled features are concatenated as inputs for the MLP architecture.

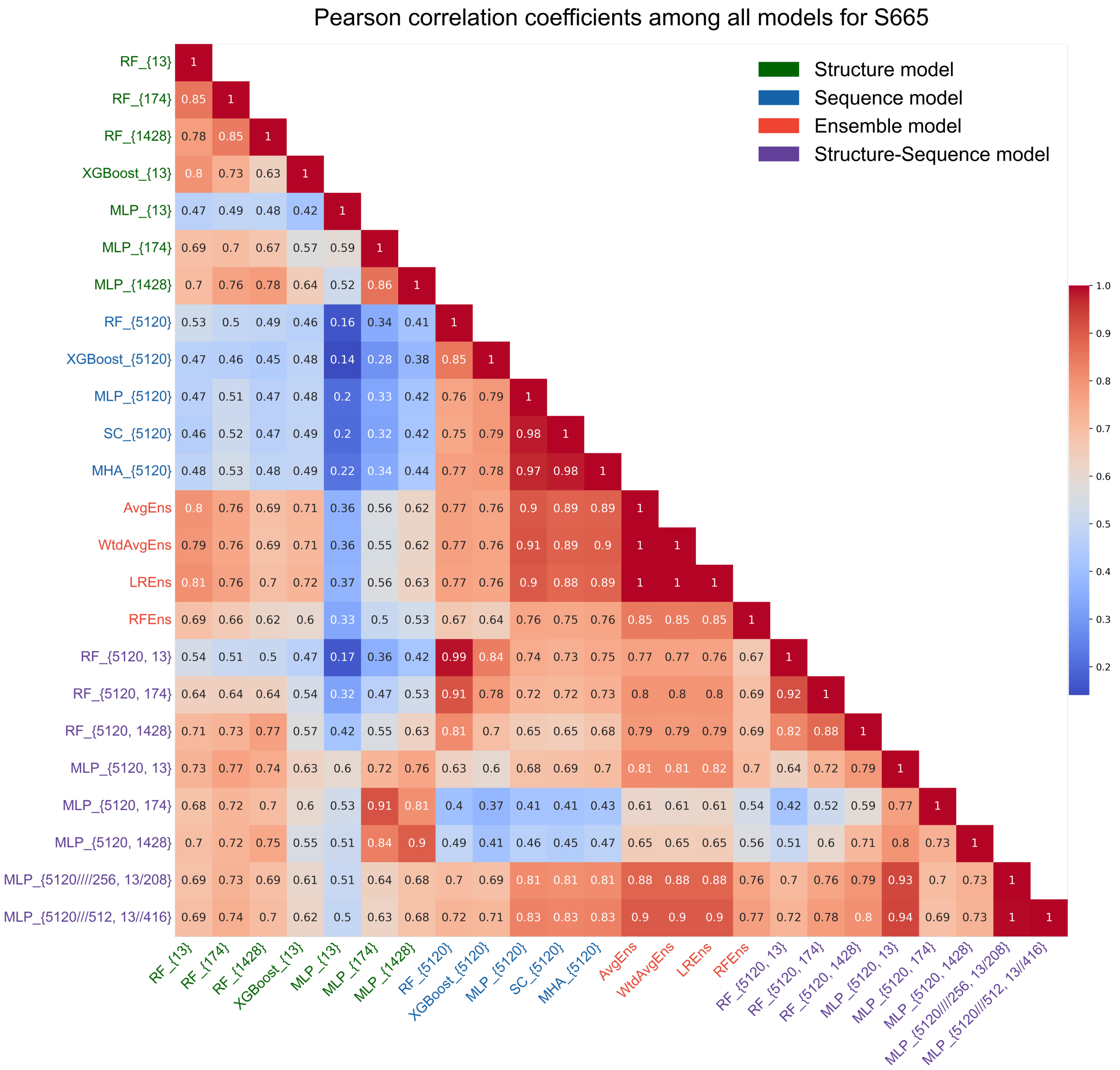


**Figure S2. Pearson correlation coefficients among all our constructed models applied to the S665 dataset.** All PCC values are statistically significantly different from zero ($P < 0.005$, *t*-test).

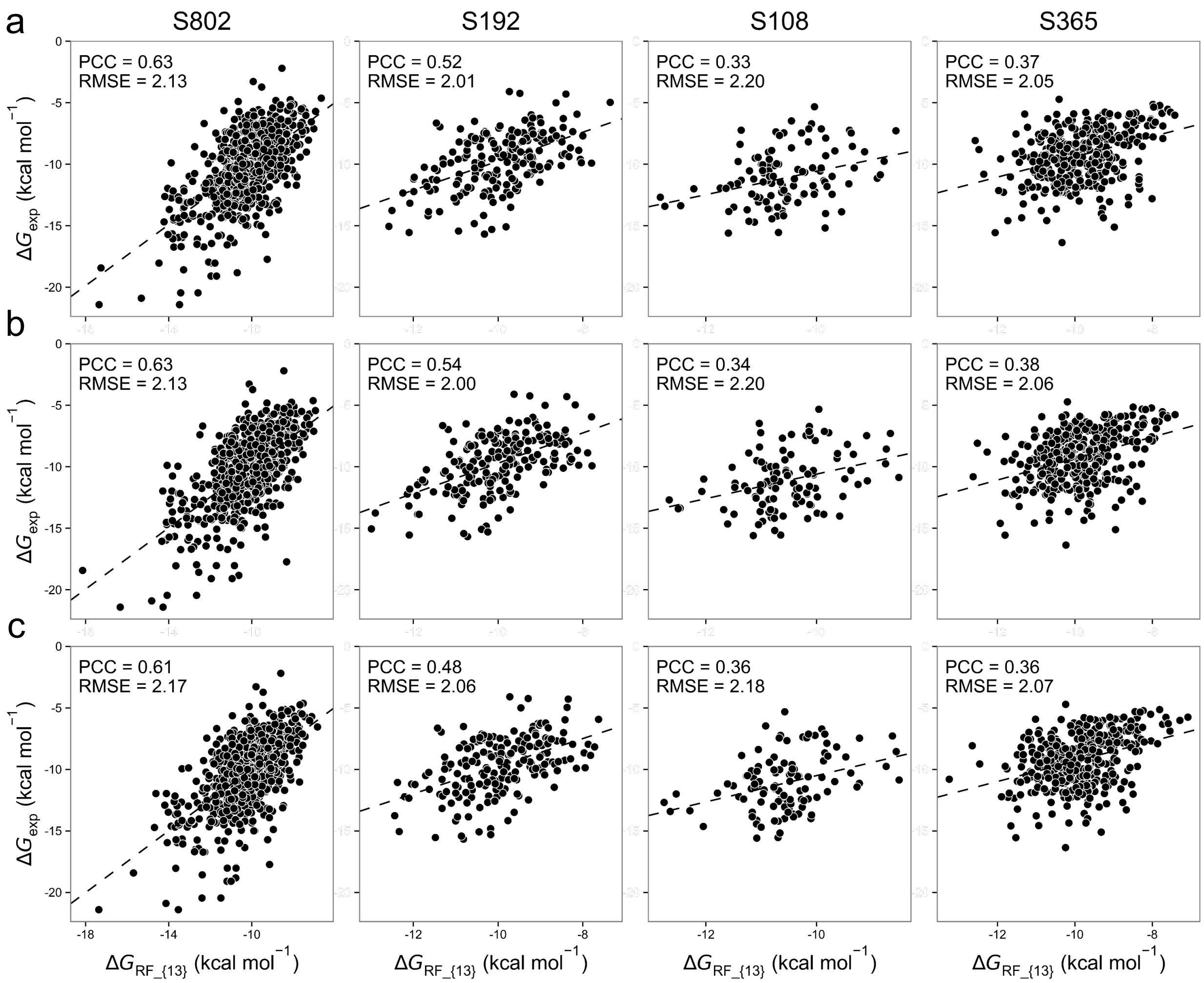


**Figure S3. Performance of RF_{13} model constructed using three different structure optimization procedures**. (a) A 100-step energy minimization with restraints on the backbone atoms (the force constant is 5 kcal mol$^{-1}$ Å$^{-2}$). (b) A 2000-step energy minimization applying the same harmonic restraints as in (a). (c) A 2000-step minimization with restraints, followed by an unconstrained 5000-step minimization. All PCC values are statistically significantly different from zero ($P < 0.005$, *t*-test).

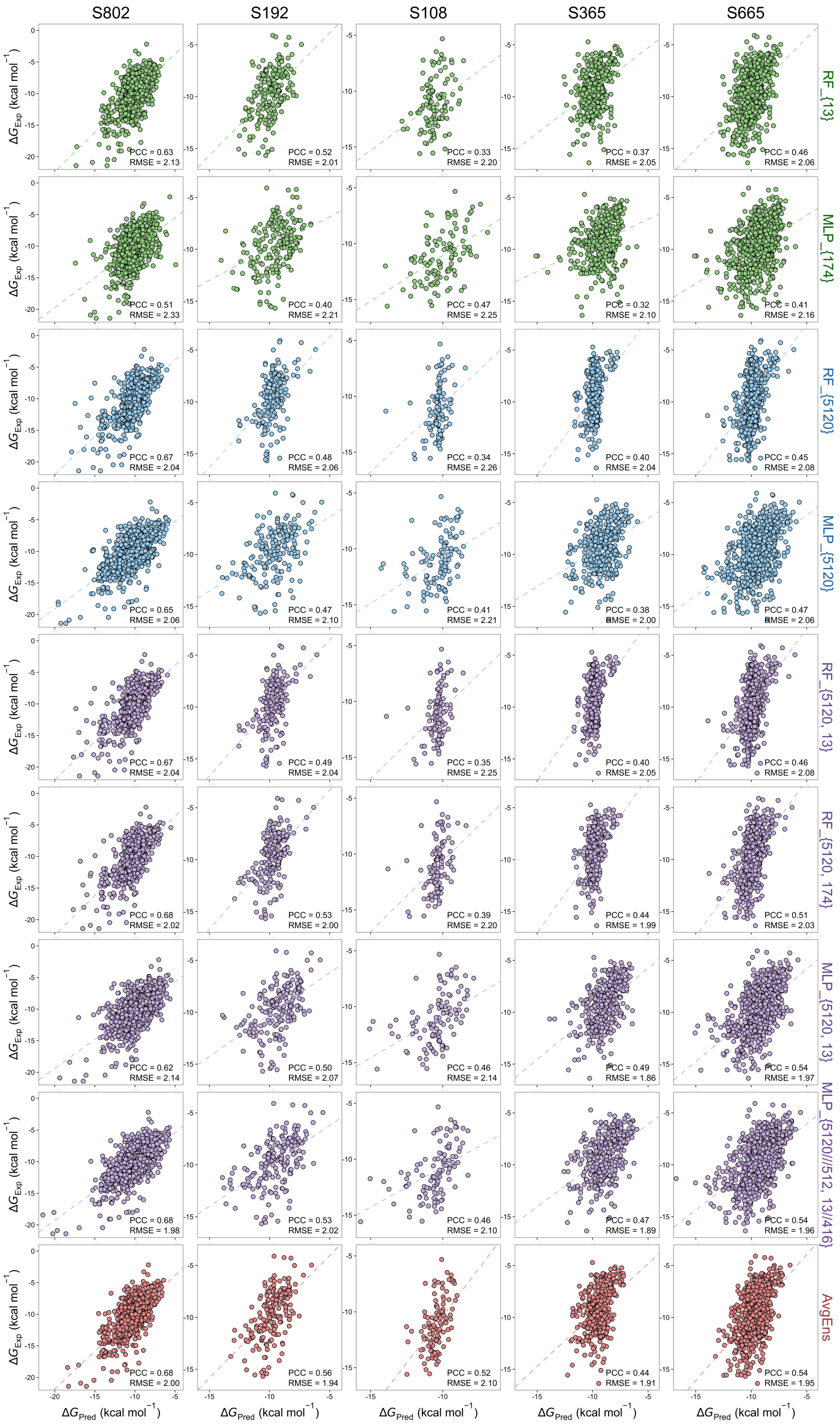


**Figure S4. Performance of nine models tested on five datasets.** 5-fold cross-validation results are shown for S802. All correlation coefficients are statistically significantly different from zero ($P < 0.005$, *t*-test). PCC: Pearson correlation coefficient, RMSE (kcal $mol^{-1}$): root-mean-square error.

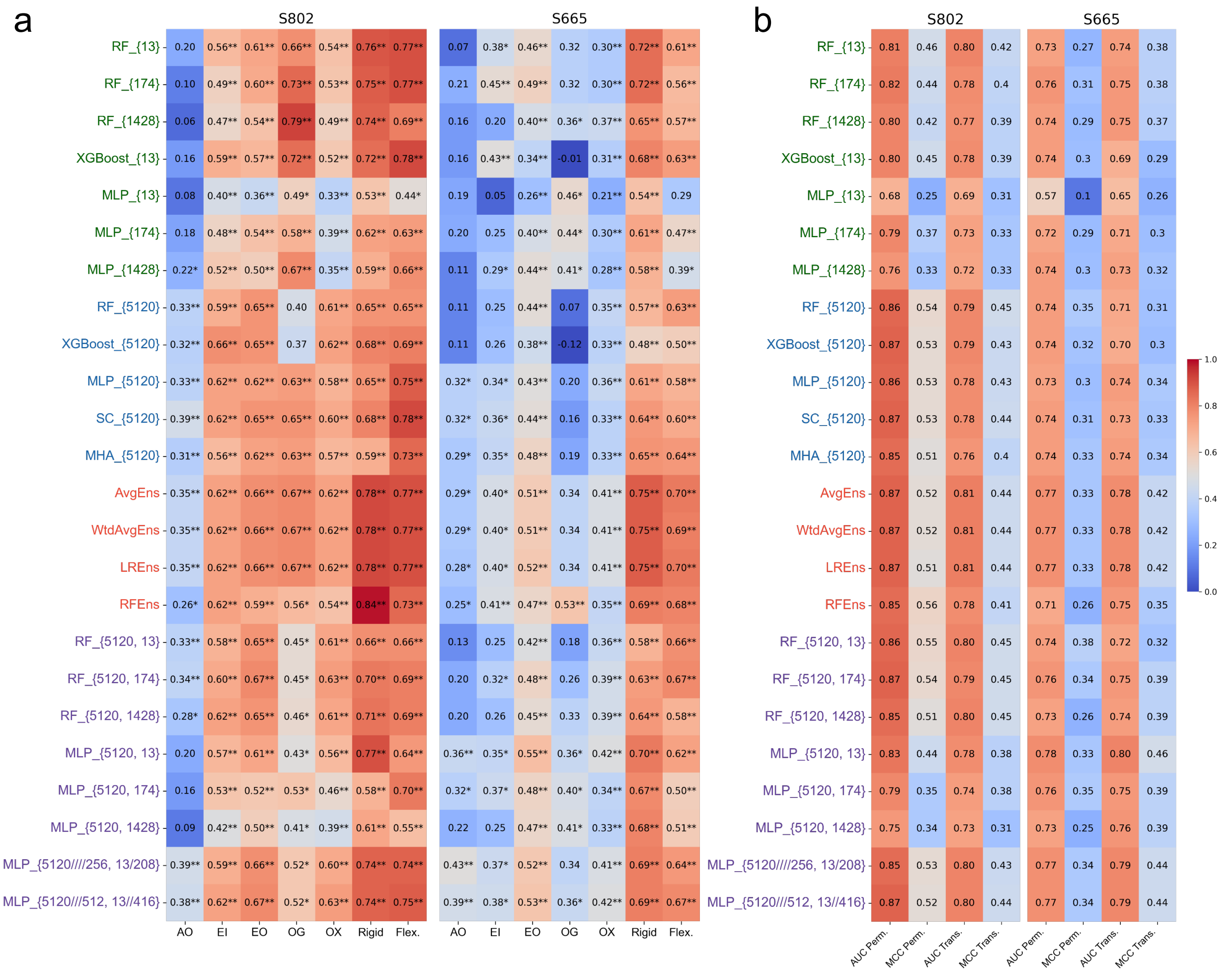


**Figure S5. Performance of all our 24 models was evaluated for three distinct interaction classifications.** (a) Pearson correlation coefficients for five functional categorizations of complexes (AO, EI, EO, OG, and OX) and rigid-body and flexible complexes. * and ** indicate statistically significant difference from zero in terms of PCC with $P < 0.05$ and $P < 0.005$ (*t*-test), respectively. (b) The values of area under the receiver operating characteristics curve (AUC) and maximal matthews correlation coefficient (MCC) values in distinguishing permanent and transient protein-protein interactions from others.

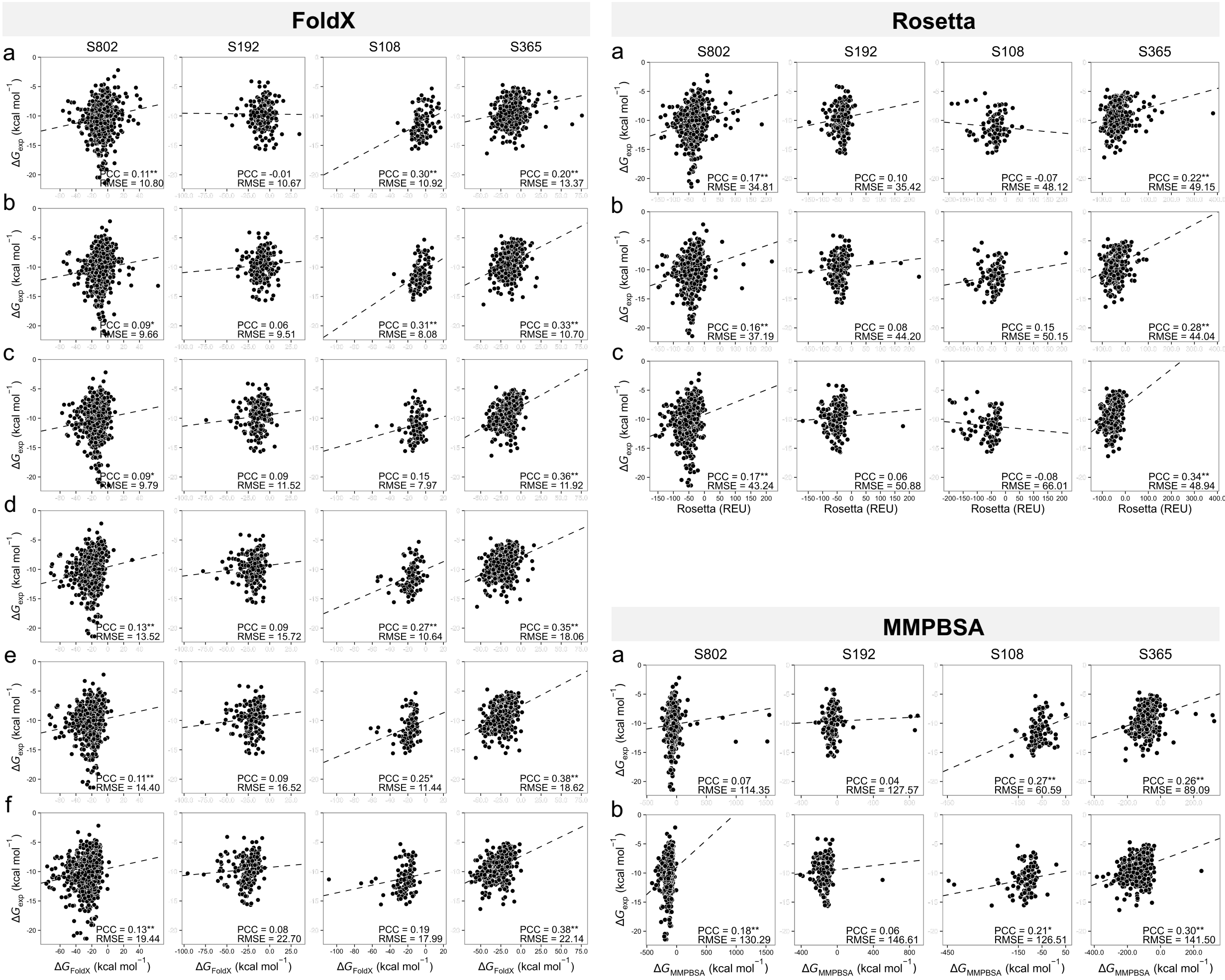


**Figure S6. Performance of FoldX, Rosetta, and MMPBSA methods under different structure optimization procedures**. For FoldX, (a), (c), and (e) present the results using experimentally-determined structures, 100-step minimized structures, and 7000-step minimized structures to perform calculation, respectively; (b), (d), and (f) provide the results involved additional optimization using the RepairPDB module to the experimentally-determined structures, 100-step minimized structures, and 7000-step minimized structures, respectively. For Rosetta, (a) experimentally determined structures, (b) 100-step minimized structures, and (c) 7000-step minimized structures were used to perform calculation. For MMPBSA, (a) 100-step minimized structures and (b) 7000-step minimized structures were used. The highest PCC values for these methods on each dataset are reported in Table S8.

**Table S1. Experimental datasets used for training and test.**

a. The number of complexes in each dataset.

| **Dataset** | **# of complexes** | **Description** |
|---|---|---|
| S802 | 802 | Training dataset, heterodimers |
| S192 | 192 | Heterodimer protein complexes |
| S108 | 108 | Heteromultimer protein complexes |
| S365 | 365 | Protein-peptide complexes |
| S665 | 665 | S192 + S108 + S365 |

b. Similarity analysis within and between datasets. The determination of the number of clusters involves the process of grouping similar complexes together based on three criteria: sequence similarity, structure similarity, and their combined use.

| **Dataset** | **Sequence** | **Structure** | **Combined** |
|---|---|---|---|
| The number of clusters in each dataset | | | |
| S802 | 525 | 539 | 582 |
| S192 | 152 | 152 | 159 |
| The number of similar complexes between S192 and S802 | | | |
| S192-S802 | 29-55 | 26-36 | 18-24 |

c. The number of complexes in each category.

| **Category** | | **S802** | **S192** | **S108** | **S365** | **S665** |
|---|---|---|---|---|---|---|
| Permanent | | 156 | 26 | 37 | 27 | 90 |
| Transient | | 177 | 58 | 15 | 120 | 193 |
| Rigid-body | | 44 | 11 | 25 | 4 | 40 |
| Flexible | | 37 | 14 | 23 | 4 | 41 |
| AO | AA | 15 | 0 | 62 | 1 | 63 |
| | AN | 64 | 4 | 0 | 1 | 5 |
| EI | | 132 | 25 | 4 | 17 | 46 |
| EO | | 264 | 70 | 17 | 108 | 195 |
| OG | | 23 | 19 | 2 | 11 | 33 |
| OX | | 304 | 74 | 23 | 227 | 324 |

**Table S2. Overview of all handcrafted features.** The table provides information on the software and references used to compute or obtain the features, as well as the number of features in each feature category.

| **Feature category** | **Number** | **Software/References** |
|---|---|---|
| Inter-protein statistical contact energies | 14 | iPot, dDFIRE, and [1-4] |
| Inter-molecular physical interaction energies, e.g., van der Waals and electrostatic | 10 | CHARMM and FoldX |
| Solvent accessible surface areas of complex and unbound partners, and the interface area | 23 | CHARMM and DSSP |
| Inter-molecular non-covalent interactions, e.g., hydrogen bonds, salt bridges, and steric classes | 66 | CHARMM, ProtInter, Arpeggio, and PRODIGY |
| Internal cavity, packing density and shape complementarity of protein-protein interface | 28 | McVol, CCP4, and [5] |
| Distribution of hot-spot and conserved residues at the interface | 21 | PROVEAN, PremPS, and FoldX |
| Sequence entropy of interface residues | 3 | [6] |
| Distribution of polar, non-polar and charged residues on the complex surface | 9 | [7] |
| Total | 174 | |
| Composition of secondary structures and residues classified according to a wide variety of properties | 1254 | AAindex database and [8] |
| Total | 1428 | |

**Table S3. Importance of each category of features for RF_{13} Model.** The importance is measured using IncNodePurity, which represents the total decrease in node impurities resulting from splitting on the variable, averaged over all trees in the random forest.

| Feature | Importance |
|---|---|
| $\boldsymbol{P_{L/SA}^{p1}}$ and $\boldsymbol{P_{L/SA}^{p2}}$ | 0.168 |
| $\boldsymbol{SA^{p1}}$ and $\boldsymbol{SA^{p2}}$ | 0.138 |
| $\boldsymbol{P_{Helix}^{IF}}$ and $\boldsymbol{P_{Sheet}^{IF}}$ | 0.117 |
| $\boldsymbol{N_{HAP}^{IF}}$ | 0.105 |
| $\boldsymbol{P_{CS}^{IF}}$ | 0.104 |
| $\Delta\boldsymbol{E_{elec}}$ | 0.096 |
| $\boldsymbol{P_{Charge}^{Surf}}$ | 0.086 |
| $\boldsymbol{S_{AI}^{IF}}$ | 0.079 |
| $\boldsymbol{E_{CE}^{IF}}$ | 0.064 |
| $\boldsymbol{V_{Cavity}}$ | 0.043 |

**Table S4. Overview of all 24 models constructed in the study.**

| Model | | Description |
|---|---|---|
| Structure-based models with handcrafted features | | |
| Traditional Machine Learning | RF_{13} | Random forest model using 13 selected handcrafted features |
| | RF_{174} | Random forest model using 174 handcrafted features |
| | RF_{1428} | Random forest model using 1428 handcrafted features |
| | XGBoost_{13} | eXtreme Gradient Boosting model using 13 selected handcrafted features |
| Deep Learning | MLP_{13} | Multilayer perceptron model using 13 selected handcrafted features |
| | MLP_{174} | Multilayer perceptron model using 174 handcrafted features |
| | MLP_{1428} | Multilayer perceptron model using 1428 handcrafted features |
| Sequence-based models with embedding features | | |
| Traditional Machine Learning | RF_{5120} | Random forest model using a 5120-dimensional embedding feature |
| | XGBoost_{5120} | eXtreme Gradient Boosting model using a 5120-dimensional embedding feature |
| Deep Learning | MLP_{5120} | Multilayer perceptron model using a 5120-dimensional embedding feature |
| | SC_{5120} | Multilayer perceptron model with skip connections using a 5120-dimensional embedding feature |
| | MHA_{5120} | Multilayer perceptron model with multi-head attention and skip connections using a 5120-dimensional embedding feature |
| Ensemble models combining structure and sequence models | | |
| Averaging Ensemble | AvgEns | Ensemble model composed of RF_{13} and MLP_{5120} using the averaging method |
| Weighted averaging Ensemble | WtdAvgEns | Ensemble model composed of RF_{13} and MLP_{5120} using the weighted averaging method |
| Linear regression | LREns | Ensemble model composed of RF_{13} and MLP_{5120} using linear regression |
| Random Forest | RFEns | Ensemble model composed of RF_{13} and MLP_{5120} using random forest |
| Structure-sequence models combining handcrafted and embedding features | | |
| Traditional Machine Learning | RF_{5120, 13} | Random forest model using a 5120-dimensional embedding feature and 13 selected handcrafted features |
| | RF_{5120, 174} | Random forest model using a 5120-dimensional embedding feature and 174 handcrafted features |
| | RF_{5120, 1428} | Random forest model using a 5120-dimensional embedding feature and 1428 handcrafted features |
| Deep Learning | MLP_{5120, 13} | Multilayer perceptron model using a 5120-dimensional embedding feature and 13 selected handcrafted features |
| | MLP_{5120, 174} | Multilayer perceptron model using a 5120-dimensional embedding feature and 174 handcrafted features |
| | MLP_{5120, 1428} | Multilayer perceptron model using a 5120-dimensional embedding feature and 1428 handcrafted features |
| | MLP_{5120////256, 13/208} | Multilayer perceptron model involves increasing the dimensionality of each handcrafted feature from 1 to 16 using one fully connected layer, and reducing the dimensionality of the 5120-dimensional complex embedding to 256 using four fully connected layers. Subsequently, the up-sampled and down-sampled features are concatenated as inputs for the MLP architecture. |
| | MLP_{5120///512, 13//416} | Multilayer perceptron model involves increasing the dimensionality of each handcrafted feature from 1 to 32 using two fully connected layers, and reducing the dimensionality of the 5120-dimensional complex embedding to 512 using three fully connected layers. Subsequently, the up-sampled and down-sampled features are concatenated as inputs for the MLP architecture. |

**Table S5. Hyperparameter search space.** The table presents a summary of the hyperparameters for each algorithm, along with the range of values that were explored during the tuning process.

| Algorithm | Tuned parameters | Initial range to sample from |
|---|---|---|
| RF | Number of Trees | [10, 20, 30, ..., 1790, 1800] |
| | Maximum Features | [auto, sqrt, log2] |
| XGBoost | Number of Trees | [10, 20, 30, 40, 50, 60, 70, 80, 90, 100, 200, 300] |
| | Maximum Depth | [1, 2, 3, 4, 5, 6, 7, 8, 9] |
| | Learning Rate | [0.00001, 0.0001, 0.001, 0.1, 0.05, 0.1, 0.15, 0.2, 0.25, 0.3, 0.35, 0.4, 0.45] |
| | Gamma | [0, 0.001, 1, 15] |
| | Subsample | [0.6, 0.7, 0.8, 0.9, 1.0] |
| | Column Sample By Tree | [0.4, 0.5, 0.6, 0.7, 0.8, 0.9, 1.0] |
| MLP | Number of Hidden Layers | [1, 2, 3, 4, 5] |
| | Learning rate | [0.000001, 0.00001, 0.0001, 0.001, 0.01] |
| | Batch Size | [8, 16, 32, 64, 128] |
| | Hidden layer 1 sizes | [16, 32, 64, 128, 256] |
| | Hidden layer 2 sizes | [8, 16, 32, 64, 128] |
| | Hidden layer 3 sizes | [4, 8, 16, 32, 64] |
| | Hidden layer 4 sizes | [4, 8, 16, 32, 64] |
| | Hidden layer 5 sizes | [2, 4, 8, 16, 32] |
| | Weight Decay | [0.0000001, 0.000001, 0.00001, 0.0001, 0.001] |

**Table S6. Number of overlapping complexes between the training sets used by other methods and the experimental datasets utilized in our study.**

| Method | Training Set | S802 | S192 | S108 | S365 |
|---|---|---|---|---|---|
| PPI-Affinity | S653 | 295 | 74 | 0 | 171 |
| PPA_Pred2 | S382 | 180 | 53 | 12 | 19 |
| Minpredictor | S139 | 68 | 20 | 32 | 8 |
| PRODIGY | S81 | 42 | 7 | 22 | 5 |
| ISLAND | S135 | 70 | 21 | 34 | 4 |

**Table S7. Performance of all models.** The best model for each subgroup is shown in bold.

| Model | S802 | | S192 | | S108 | | S365 | | S665 | |
|---|---|---|---|---|---|---|---|---|---|---|
| | PCC | RMSE | PCC | RMSE | PCC | RMSE | PCC | RMSE | PCC | RMSE |
| Structure-based models with handcrafted features | | | | | | | | | | |
| **RF_{13}** | **0.63** | **2.13** | **0.52** | **2.01** | **0.33** | **2.20** | **0.37** | **2.05** | **0.46** | **2.06** |
| RF_{174} | 0.61 | 2.17 | 0.54 | 1.98 | 0.35 | 2.17 | 0.40 | 1.95 | 0.49 | 2.00 |
| RF_{1428} | 0.58** | 2.22 | 0.51 | 2.03 | 0.41 | 2.08 | 0.39 | 2.02 | 0.48 | 2.03 |
| XGBoost_{13} | 0.62 | 2.14 | 0.51 | 2.01 | 0.37 | 2.31 | 0.22** | 2.18 | 0.41* | 2.16 |
| **MLP_{174}** | **0.51** | **2.33** | **0.40** | **2.21** | **0.47** | **2.25** | **0.32** | **2.10** | **0.41** | **2.16** |
| MLP_{13} | 0.34** | 2.57 | 0.19** | 2.40 | 0.28* | 2.58 | 0.22 | 2.09 | 0.23** | 2.27 |
| MLP_{1428} | 0.51 | 2.34 | 0.44 | 2.13 | 0.41 | 2.21 | 0.32 | 2.14 | 0.44 | 2.15 |
| Sequence-based models with embedding features | | | | | | | | | | |
| **RF_{5120}** | **0.67** | **2.04** | **0.48** | **2.06** | **0.34** | **2.26** | **0.40** | **2.04** | **0.45** | **2.08** |
| XGBoost_{5120} | 0.69* | 1.97 | 0.48 | 2.04 | 0.27 | 2.31 | 0.31** | 2.02 | 0.43 | 2.08 |
| **MLP_{5120}** | **0.65** | **2.06** | **0.47** | **2.10** | **0.41** | **2.21** | **0.38** | **2.00** | **0.47** | **2.06** |
| SC_{5120} | 0.67** | 2.01 | 0.47 | 2.10 | 0.38 | 2.30 | 0.36* | 2.06 | 0.46 | 2.11 |
| MHA_{5120} | 0.63* | 2.12 | 0.47 | 2.11 | 0.38 | 2.42 | 0.38 | 2.03 | 0.47 | 2.12 |
| Ensemble models combining structure and sequence models | | | | | | | | | | |
| **AvgEns** | **0.68** | **2.00** | **0.56** | **1.94** | **0.52** | **2.10** | **0.44** | **1.91** | **0.54** | **1.95** |
| WtdAvgEns | 0.68 | 2.00 | 0.56 | 1.94 | 0.52 | 2.10 | 0.44 | 1.91 | 0.54 | 1.95 |
| LREns | 0.68 | 1.97 | 0.56 | 1.92 | 0.52 | 2.09 | 0.44 | 1.90 | 0.54 | 1.94 |
| RFEns | 0.63** | 2.12 | 0.53 | 2.01 | 0.46 | 2.18 | 0.39 | 1.99 | 0.49* | 2.03 |
| Structure-sequence models combining handcrafted and embedding features | | | | | | | | | | |
| **RF_{5120, 174}** | **0.68** | **2.02** | **0.53** | **2.00** | **0.39** | **2.20** | **0.44** | **1.99** | **0.51** | **2.03** |
| RF_{5120, 13} | 0.67* | 2.04 | 0.49 | 2.04 | 0.35 | 2.25 | 0.40 | 2.05 | 0.46** | 2.08 |
| RF_{5120, 1428} | 0.68 | 2.02 | 0.55 | 1.97 | 0.31 | 2.27 | 0.42 | 2.01 | 0.49 | 2.04 |
| **MLP_{5120///512, 13//416}** | **0.68** | **1.98** | **0.53** | **2.02** | **0.46** | **2.10** | **0.47** | **1.89** | **0.54** | **1.96** |
| MLP_{5120, 13} | 0.62** | 2.14 | 0.50 | 2.07 | 0.46 | 2.14 | 0.49 | 1.86 | 0.54 | 1.97 |
| MLP_{5120, 174} | 0.54** | 2.39 | 0.48 | 2.24 | 0.48 | 2.09 | 0.36* | 2.25 | 0.47** | 2.22 |
| MLP_{5120, 1428} | 0.49** | 2.46 | 0.49 | 2.11 | 0.49 | 2.02 | 0.36** | 2.28 | 0.45** | 2.19 |
| MLP_{5120////256, 13/208} | 0.66** | 2.03 | 0.52 | 2.02 | 0.46 | 2.11 | 0.46* | 1.90 | 0.54** | 1.97 |

PCC: Pearson correlation coefficient between experimental and predicted binding affinities. RMSE (kcal $mol^{-1}$): root-mean-square error. All presented values of correlation coefficients are statistically significantly different from zero ($P < 0.05$, *t*-test). *$P < 0.05$/ **$P < 0.005$ compared to the best model in each subgroup (Hittner2003 test).

**Table S8. Comparison of methods' performances.** Our three models outperform all the other approaches in terms of PCC with $P < 0.05$ (Hittner2003 test), except for the PCC values with superscripts of 'a', 'm', or 'r', indicating that these models do not have statistically significant differences from AvgEns, MLP_{5120}, or RF_{13}, respectively.

| Method | S802 | | S192 | | S108 | | S365 | |
|---|---|---|---|---|---|---|---|---|
| | PCC | RMSE | PCC | RMSE | PCC | RMSE | PCC | RMSE |
| AvgEns | 0.68 | 2.00 | 0.56 | 1.94 | 0.52 | 2.10 | 0.44 | 1.91 |
| MLP_{5120} | 0.65 | 2.06 | 0.47 | 2.10 | 0.41 | 2.21 | 0.38 | 2.00 |
| RF_{13} | 0.63 | 2.13 | 0.52 | 2.01 | 0.33 | 2.20 | 0.37 | 2.05 |
| PPI-Affinity | 0.42 | 2.49 | $0.34^{m}$ | 2.24 | NA | NA | 0.22 | 3.89 |
| PPA_Pred2 | 0.34 | 3.00 | $0.30^{m}$ | 2.55 | NA | NA | - | 3.20 |
| Minpredictor | 0.31 | 2.76 | 0.25 | 2.84 | NA | NA | 0.20 | 2.52 |
| PRODIGY | 0.29 | 3.14 | 0.21 | 3.61 | $0.26^{mr}$ | 2.81 | $0.30^{mr}$ | 2.93 |
| ISLAND | 0.32 | 2.58 | 0.15 | 2.39 | NA | NA | 0.26 | 2.24 |
| FoldX | 0.11 | 14.40 | - | 16.52 | $0.25^{mr}$ | 11.44 | $0.38^{amr}$ | 18.62 |
| MMPBSA | - | 114.35 | - | 127.57 | $0.27^{mr}$ | 60.59 | $0.26^{mr}$ | 89.09 |
| Rosetta | 0.16 | 37.19 | - | 44.20 | - | 50.15 | $0.28^{mr}$ | 44.04 |
| ZRANK | 0.14 | 109.43 | - | 122.67 | $0.20^{r}$ | 101.59 | 0.25 | 123.61 |
| ZRANK2 | 0.12 | 320.10 | - | 334.78 | $0.22^{mr}$ | 300.86 | 0.13 | 366.46 |
| RosettaDock | 0.15 | 10.25 | - | 16.40 | $0.38^{amr}$ | 6.08 | - | 18.08 |
| pyDock | 0.23 | 35.22 | 0.16 | 39.95 | $0.32^{mr}$ | 30.39 | $0.28^{mr}$ | 44.47 |
| SIPPER | -0.10 | 18.64 | -0.17 | 18.60 | -0.19 | 19.24 | -0.24 | 21.76 |
| PISA | 0.24 | 10.18 | - | 9.52 | - | 10.98 | $0.34^{amr}$ | 9.03 |
| FireDock | 0.13 | 67.30 | 0.15 | 77.83 | $0.31^{mr}$ | 52.93 | $0.25^{m}$ | 90.38 |
| FireDock_AB | 0.12 | 85.08 | - | 96.89 | $0.29^{mr}$ | 71.46 | 0.23 | 105.32 |
| FireDock_EI | 0.18 | 33.87 | 0.20 | 39.51 | $0.37^{amr}$ | 24.06 | $0.27^{mr}$ | 46.45 |
| PIE | -0.17 | 12.18 | - | 11.70 | -0.25 | 12.92 | -0.36 | 11.37 |

PCC: Pearson correlation coefficient between experimental and predicted binding affinities. RMSE (kcal $mol^{-1}$): root-mean-square error. Only correlation coefficients statistically significantly different from zero are shown ($P < 0.05$, *t*-test). '-' indicates that the PCC value is not statistically significantly different from zero. 'NA': not available, these four methods cannot calculate binding affinity for multimers.
